# Formin-Dependent Adhesions are Required for Invasion by Epithelial Tissues

**DOI:** 10.1101/124297

**Authors:** Tim B. Fessenden, Yvonne Beckham, Mathew Perez-Neut, Aparajita H. Chourasia, Kay F. Macleod, Patrick W. Oakes, Margaret L. Gardel

## Abstract

Developing tissues change shape, and tumors initiate spreading, through collective cell motility. Conserved mechanisms by which tissues initiate motility into their surroundings are not known. We investigated cytoskeletal regulators during collective invasion by mouse tumor organoids and epithelial MDCK acini undergoing branching morphogenesis. Inhibition of formins, but not Arp2/3, prevented the formation of migrating cell fronts in both cell types. MDCK cells depleted of the formin protein Dia1 formed polarized acini and could execute planar cell motility, either within the acinus or in 2D scattering assays. However, Dia1 was required to form protrusions into the collagen matrix. Live imaging of actin, myosin, and collagen in control acini revealed adhesions that deformed individual collagen fibrils, while Dia1-depleted acini exhibited unstable adhesions with minimal collagen deformation. This work identifies Dia1-mediated adhesions as essential regulators of tissue shape changes, through their role in focal adhesion maturation.

## Introduction

Tissue shape changes encompass multiple developmental and pathological processes. In order to form branched tubular networks, developing tissues such as mammalian vasculature or the Drosophila trachea undergo extensive elongation and remodeling known as branching morphogenesis (Lecaudey and Gilmour, 2006; Lubarsky and Krasnow, 2003; Wang et al., 2017). In many cases, branching morphogenesis is initiated when growth factors stimulate a few individual cells within the developing tissue to extend protrusions that adhere to the surrounding extracellular matrix (ECM). These cells subsequently lead cohorts of their neighbors out of their initial site, migrating collectively through the ECM to form extensively branched tubules (Affolter et al., 2009; O’Brien et al., 2002). Malignant tissue can exhibit similar, if deregulated, shape changes during local invasion from the site of tumor formation (Friedl et al., 2012). Invasion by tumors is often accomplished by collective cell migration, in a manner that frequently mimics development (Friedl and Alexander, 2011; Gray et al., 2010). In both developmental and pathological contexts, shape changes undertaken by tissues rely on the coordination of cell motility and cell adhesions to neighboring cells and the ECM.

An outstanding question is how tissues transition from compact structures dominated by cell-cell adhesions to invading cohorts of cells that interact extensively with their ECM. A well-established framework describing the acquisition of invasive behaviors is the epithelial-mesenchymal transition (EMT) (Thiery et al., 2009). EMT comprises a gene regulatory program that simultaneously suppresses cells’ epithelial traits while activating mesenchymal traits, thereby stimulating invasion. However, EMT does not adequately describe tissue shape changes when epithelial traits such as cell-cell adhesion are maintained (Affolter et al., 2009; Kowalski et al., 2003; Shamir et al., 2014). In these cases, a partial or transient EMT has been proffered to account for invasive behaviors exhibited by intact tissues (Christiansen, 2006; Friedl et al., 2012; Lambert et al., 2017; O’Brien et al., 2002; Revenu and Gilmour, 2009). But this model leaves unclear how the partial loss or gain of epithelial or mesenchymal traits, respectively, can orchestrate collective cell invasion (Ewald et al., 2012; O’Brien et al., 2004). For example, cell movements within tissues are required in some cases to maintain epithelial homeostasis, (Haigo and Bilder, 2011; Isabella and Horne-Badovinac, 2016; Wang et al., 2013) but in other cases to drive branching morphogenesis (Ewald et al., 2008; Wang et al., 2017). Thus we lack precise mechanisms to describe how motility and adhesions to the ECM are shifted in individual cells to accomplish tissue shape changes.

Cell motility and adhesions rely on the actin cytoskeleton, which is organized in space and time into protrusive, contractile and adhesive organelles (Lauffenburger and Horwitz, 1996). Protrusion of the cell’s leading edge is typically driven by Arp2/3–mediated lamellipodia (Gardel et al., 2010; Pollard and Borisy, 2003). Proximal to the lamellipodia, within a RhoA-dependent lamella, actomyosin networks construct actin bundles and generate contractile forces. Coordinated with the actin cytoskeleton is the assembly and modification of focal adhesions, which serve as sites of biochemical signaling and as mechanical linkages between the cell and its surroundings (Gardel et al., 2010; Geiger and Yamada, 2011). Focal adhesions assemble within the lamellipodia (Zaidel-Bar et al., 2003), but undergo increases in size and changes in composition in a ‘maturation’ process that relies on Rho effectors Myosin II (Riveline et al., 2001) and Dia1 (Chrzanowska-Wodnicka and Burridge, 1996; Oakes et al., 2012). Focal adhesion maturation has been extensively studied in cells on 2D planar surfaces, and exerts context-dependent effects on matrix deposition, front-rear polarity, and migration speed (Hoffman et al., 2006; Horton et al., 2016; Oakes et al., 2012; Rahman et al., 2016; Thievessen et al., 2013). In fibrillar 3D environments, focal adhesion morphology is significantly altered and the role of focal adhesion maturation is less well-defined (Doyle et al., 2015; Fraley et al., 2010; Harunaga and Yamada, 2011; Kubow et al., 2013). While branching morphogenesis and tumor invasion require canonical focal adhesion components, (Friedl et al., 2012; Hunter and Zegers, 2010; Jiang et al., 2001; Wei et al., 2009) the mechanism underlying their regulation is not known.

We used 3D culture and organotypic models to dissect the contributions of cytoskeletal organelles to tissue shape changes. A tractable *in vitro* model to study the regulation of tissue remodeling is provided by Manin Darby Canine Kidney (MDCK) cells undergoing branching morphogenesis. MDCK cells cultured in 3D matrices form hollow acini that resemble simple cuboidal epithelial tissues (Bryant and Mostov, 2008). These undergo robust and well-described branching morphogenesis in response to Hepatocyte Growth Factor (HGF) (Yu et al., 2003). Using pharmacological inhibitors, we found that branching morphogenesis requires the activity of formins but not Arp2/3. Tumor explants from mice confirmed that formins were also required for multicellular invasion. We found that the formin isoform Dia1 was dispensable for growth and polarization of MDCK acini and, interestingly, Dia1 depletion did not affect HGF-mediated planar cell motility in either cell scattering assays or cell motility within acini. Rather, Dia1 was selectively required for cells to form stable, mature adhesions and deform collagen fibrils during the initiation of branching morphogenesis. Interestingly, we found that adhesion sites to collagen fibrils contained both actin and myosin II, whose accumulation coincided with force generation on collagen fibrils. Dia1-depleted cells exhibited unstable myosin-rich adhesions, coincident with the loss of dense puncta and linear bundles of actin at the basal cortex. Thus, Dia1 conditions tissue shape changes by controlling the stabilization of cell-collagen adhesions.

## Results

### Formin activity is required for invasion and branching morphogenesis

To explore the roles of Arp2/3 and formins in tissue shape changes, we used two cell models:Manin-Darby Canine Kidney (MDCK) acini undergoing branching morphogenesis, and invasive motility by murine tumor explants. Single MDCK cells embedded in Matrigel grew into polarized acini with clear lumens over 4-5 days. Using a protocol modified from Rubashkin et al, we isolated acini and plated them intact into 2 mg/ml collagen gels (Rubashkin et al., 2014). Once plated in collagen, acini remained quiescent for at least 1 week or could be induced to undergo branching morphogenesis by the addition of 20 ng/ml Hepatocyte Growth Factor (HGF) (Fig. 1A and Materials and Methods). After 48 hours, branching morphogenesis resulted in multiple protrusive fronts extending into the collagen gel from each acinus. These protrusions started as extensions by single cells which then lengthened to form chains and tubules of cells, as shown by phalloidin staining for actin (Fig S1). The size and growth rate of protrusions we observed agreed with the stages of branching morphogenesis described previously (Yu et al., 2003; Zegers et al., 2003).

**Figure 1:**
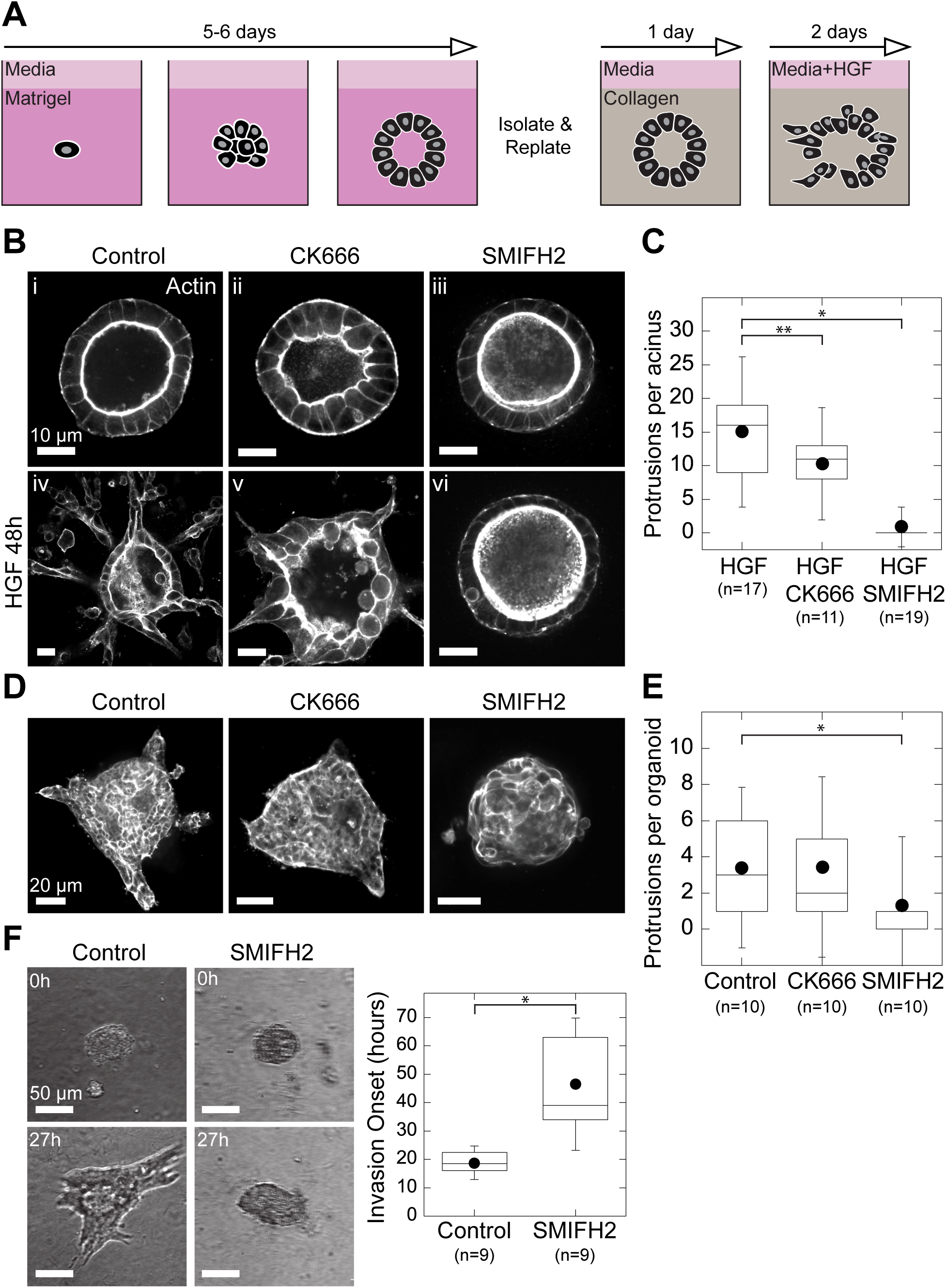
Formin Activity is Required for Invasion and Branching Morphogenesis. **A,** Strategy for 3D culture and branching morphogenesis of Manin Darby Canine Kidney (MDCK) acini. **Bi-iii,** Equatorial confocal sections of MDCK acini showing phalloidin stain for f-actin. Acini were plated in collagen and treated with 50 μM CK666 or 30 μM SMIFH2 for 48 hrs. **Biv-vi**, F-actin in acini treated for 48 hrs with 20ng/ml Hepatocyte Growth Factor (HGF) with or without the indicated inhibitors. **C,** Box plot of protrusions formed per acinus in each condition. **D,** Mouse mammary tumors from MMTV/PyMT mice. Tumors were harvested and digested, and resulting organoids were plated directly in collagen gels and treated with serum alone or with 50 μM CK666 or 30 μM SMIFH2, then fixed after 24 hrs. Shown are equatorial confocal sections of phalloidin stain for f-actin. **E,** Box plot of protrusions per organoid, with number of organoids scored indicated below. **F**, Organoids were plated in collagen, treated with serum alone or with 30 μM SMIFH2, and imaged in brightfield for 72 hrs by time lapse microscopy (see also Video 1). Invasion onset was scored as the time of initial extension into collagen gel and plotted in box plot with number of organoids scored indicated below. Box plots show the 25^th^ and 75^th^ percentiles and the median, circle indicates mean, and whiskers mark 1.5 standard deviations. Single asterisk indicates p<0.01, double asterisk indicates p<0.05, by a Student's two tailed t-test assuming unequal variance.

To explore the role of Arp2/3 and formins in regulating branching morphogenesis, we treated acini with the Arp2/3 inhibitor CK666 (Nolen et al., 2009) or the pan-formin inhibitor SMIFH2 (Rizvi et al., 2009). Phalloidin staining in control acini, prior to branching morphogenesis, revealed that F-actin was localized exclusively to the cell membranes. The apical surface was marked by a dense band of actin, due to the presence of microvilli as previously reported (McAteer et al., 1987) (Fig. 1B, i). Addition of 20ng/ml HGF resulted in prototypical branching morphogenesis (Fig. 1B, iv). Treatment with 50 μM CK666 in the absence of HGF caused individual cells to bulge within acini, as evidenced by convex curvature at the apical membrane (Fig. 1B, ii). This suggested an acute polarity defect, consistent with reported roles for Arp2/3 in forming polarized membrane domains (Martin-Belmonte et al., 2007). However, stimulating acini with HGF in the presence of CK666 did not prevent acini from forming multiple protrusions into the collagen gel even as cell morphology remained perturbed (Fig 1B, v). In contrast to Arp2/3 inhibition, treating acini with 30 μM SMIFH2 did not appreciably alter acinar morphology relative to controls (Fig. 1B, iii). When stimulated HGF, however, SMIFH2-treated acini did not form any discernable protrusions into the collagen gel. Rather, these acini were indistinguishable from untreated controls (Fig. 1B, vi).

We scored the ability of cells to respond to HGF in the above conditions by counting protrusive structures, whether by single cells or multicellular chains or branches invading into the collagen gel. This confirmed that SMIFH2, but not CK666, prevented protrusion formation in response to HGF (Fig. 1C). Together, these data demonstrate that Arp2/3 activity is dispensable for invasion during branching morphogenesis, despite its role in polarity signaling. Meanwhile, formin activity is dispensable for acinar morphology, but required for formation of HGF-mediated invasion into the surrounding collagen matrix.

To confirm the generality of these results, we performed invasion assays with primary tumor organoids from mice. Tumors from the Murine Mammary Tumor Virus-Polyoma Middle-T (MMTV-PyMT) mouse strain comprise heterogeneous cell populations, do not form a basement membrane, and do not require a defined signal to stimulate invasion into collagen gels (Cheung et al., 2013; Lin et al., 2003). Mice with advanced tumors were sacrificed and tumor tissue was harvested and digested into multicellular organoids. Organoids were first cultured in Matrigel for 48 hours in low-serum media, then replated intact into collagen gels. To stimulate invasion, media containing serum was added and after 24 hours tumor organoids were fixed to visualize the actin cytoskeleton. Control organoids were uniformly invasive, extending multicellular fronts into the collagen matrix (Fig. 1D). Inhibition of Arp2/3 did not affect the total number of fronts per organoid. However, SMIFH2-treated organoids did not form invasive fronts after 24 hours (Fig. 1E). To clarify the motility defects rendered by formin inhibition, we analyzed invasion in tumor organoids via time-lapse imaging (Video 1). Cells in control and SMIFH2-treated tumor organoids appeared to actively rearrange over 48 hours. However, by 20 hours after plating, control organoids had formed subcellular extensions from which collective invasion proceeded. Formin inhibition prevented organoids from generating such extensions until ∼40 hours following plating (Fig. 1F).

Collectively, these data implicate formin activity as a previously unappreciated determinant of both branching morphogenesis by MDCK acini and collective invasion by mouse tumor organoids. In MDCK acini, formin activity appears to be dispensable for cell shape and acinar homeostasis prior to HGF stimulation. However, in both acini and tumor organoids, formin inhibition blocked tissue shape changes by preventing the formation of the invasive fronts into the surrounding collagen matrix.

### Dia1 and FHOD1 are required for branching morphogenesis

Because SMIFH2 is a pleiotropic inhibitor of formins (Rizvi et al., 2009), we sought to identify formin family members required for invasion. We initially chose to explore the roles of three formin family members: Diaphanous 1 (Dia1), Diaphanous 2 (Dia2), and Formin Homology 2 Domain Containing 1 (FHOD1). We and others have previously established roles for Dia1 in cell motility and focal adhesion maturation (Oakes et al., 2012; Riveline et al., 2001). Like Dia1, Dia2 is activated by Rho and participates in stress fiber formation but is also implicated in cytokinetic ring formation (Gupton et al., 2007; Pollard, 2010). FHOD1 is a noncanonical formin activated through Rac that has been shown to bind ROCK and participates in focal adhesion maturation (Iskratsch et al., 2013; Takeya et al., 2008). shRNA constructs targeting these genes reduced the expression to 45-50% of control levels (Fig. S2A).

Stable cell lines expressing each of these shRNA constructs could proliferate in 2D culture and in Matrigel. After replating acini from each into collagen gels, we fixed and stained them to analyze acinar morphology and actin architecture. While shDia1 and shFHOD1 cells could form acini with cleared lumens, shDia2 cells formed acini of aberrant shapes without clear lumens (Fig. 2A, top row and Fig. S2B).

**Figure 2:**
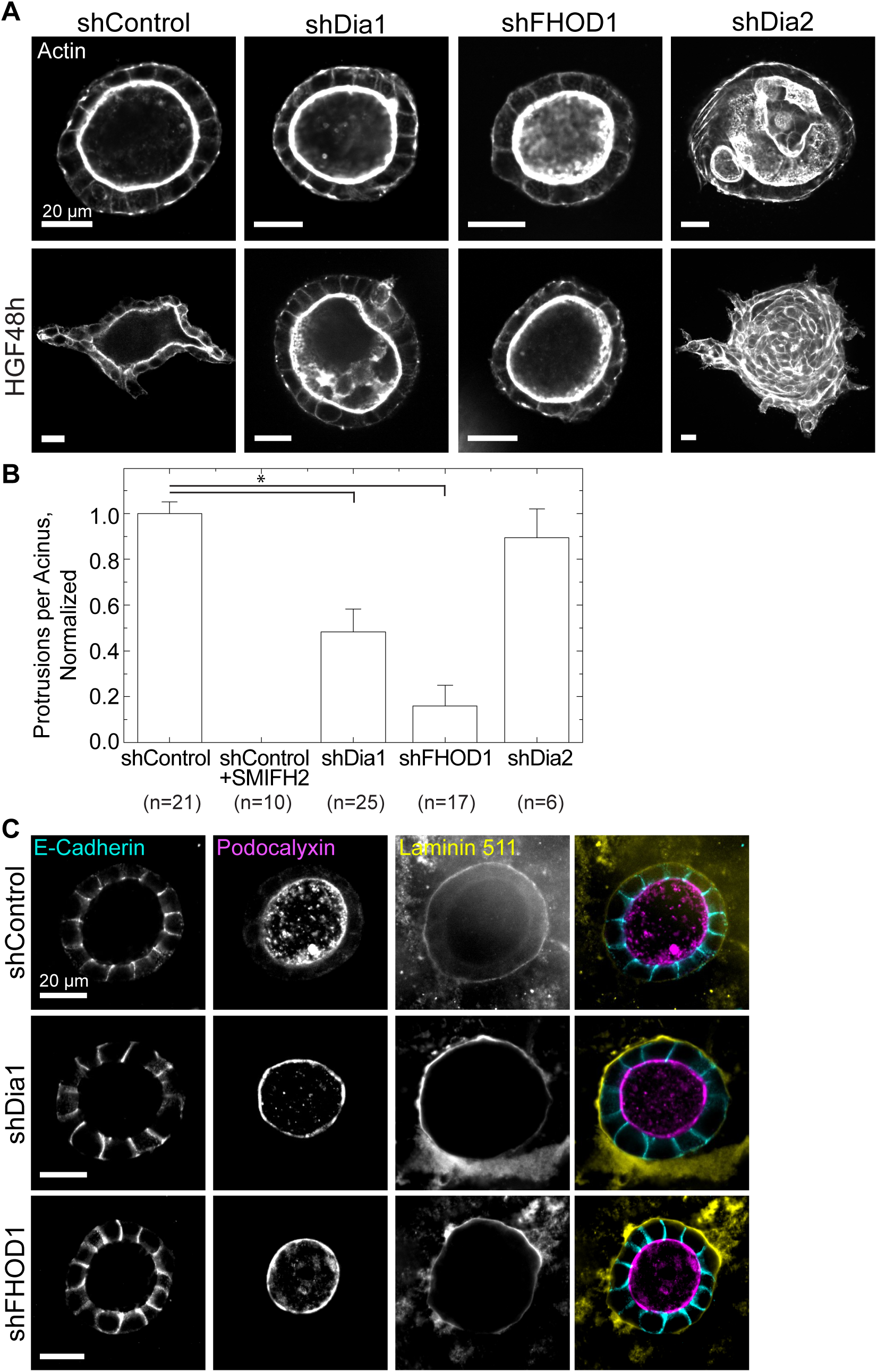
Dia1 and FHOD1 are Required for Branching Morphogenesis. **A,** Equatorial confocal sections of actin stain in acini of each genotype prior to (top) and following stimulation with HGF for 48 hrs (bottom). **B,** The number of protrusions per acinus relative to matched shControl acini; number of acini scored indicated below. SMIFH2 treatment serves as a negative control. See Fig. S2. **C,** Equatorial confocal sections of acini stained for E-cadherin to mark cellcell junctions, podocalyxin to mark the apical surface, and the basement membrane component Laminin 511 in acini generated from control (top), shDia1 (middle) and shFHOD1 (bottom) cells. Asterisk indicates p<0.01 by a Student's two tailed t-test assuming unequal variance.

To test their capacity to undergo branching morphogenesis, we treated acini from each cell type with 20 ng/ml HGF. After 48 hours, shDia1 and shFHOD1 acini formed significantly fewer invasive protrusions relative to shControl acini (Fig. 2A, bottom row). We analyzed protrusions as above and compared the average number of protrusions formed per acinus in each genotype (Fig. S2C). Combining data from all acini showed a significant decrease in protrusions formed by shDia1 and shFHOD1 acini, but not shDia2 acini, relative to matched controls (Fig. 2B). When protrusion lengths were measured across these conditions, we found that the protrusions formed by shDia1 and shFHOD1 acini were shorter on average than matched controls (Fig. S2D).

Although they appeared morphologically similar to controls, knockdown of Dia1 and FHOD1 may still alter cell polarity within the acini. To test for polarity defects in shDia1 and shFHOD1 acini we stained for E-cadherin, Podocalyxin, and Laminin 511 by immunofluorescence. Localization of these proteins to lateral, apical, and basal surfaces, respectively, resembled that in control acini (Fig. 2C). These results suggest that as they develop and polarize, shDia1 and shFHOD1 acini could generate and orient an apical domain, marked by podocalyxin, and form adherens junctions, marked by E-cadherin. After replating in collagen, these acini also deposited basement membrane laminins similar to their control counterparts.

We conclude from these data that depletion of Dia1 and FHOD1 does not impair the growth and polarization of acini. These formins are, however, required to initiate branching morphogenesis by establishing and elongating protrusions into the collagen gel. In contrast, Dia2 is required for normal acinar development but is dispensable for generating invasive protrusions in response to HGF. Dia1 and FHOD1 appear to play nonoverlapping roles, perhaps due to their differing upstream activators. We were interested in the role played by Dia1 because another RhoA effector, ROCK, is known to restrict rather than promote protrusions in response to HGF (Yu et al., 2003). This suggests that these two RhoA effectors may play competing roles in controlling branching morphogenesis. We therefore sought to clarify the role of Dia1.

### Dia1 is dispensable for HGF-mediated planar motility in 2D and within acini

Branching morphogenesis requires that cells migrate into the surrounding collagen matrix (Yu et al., 2003). Previous work has shown the importance of Dia1 in establishment of a polarized leading edge for motility on 2D substrates (Dachsel et al., 2013; Isogai et al., 2015). Thus, one possibility is that a cell migration defect underlies the role of Dia1 in branching morphogenesis.

To test this hypothesis, we first used a cell scattering assay, in which HGF drives the dissociation of cell islands on glass coverslips into individual, highly motile cells (Stoker et al., 1987). We plated shControl and shDia1 MDCK cells on glass coverslips and serum starved them for 24 hours before adding 20 ng/ml HGF to stimulate scattering. Timelapse microscopy revealed that shDia1 cells could break cell-cell contacts and migrate as single cells similarly to shControl cells (Fig. 3A and Video 2). While scattering occurred in both cell types, colonies of shDia1 cells took ∼1 hour longer to begin scattering (Fig. S2A and B). We tracked individual cells (Fig. 3B) and found a modest reduction in the mean instantaneous speed of shDia1 cells, but no change in their persistence (Fig. 3C). Thus, Dia1 may contribute to the ability to initiate scattering, but is dispensable for HGF-mediated motility in 2D.

**Figure 3:**
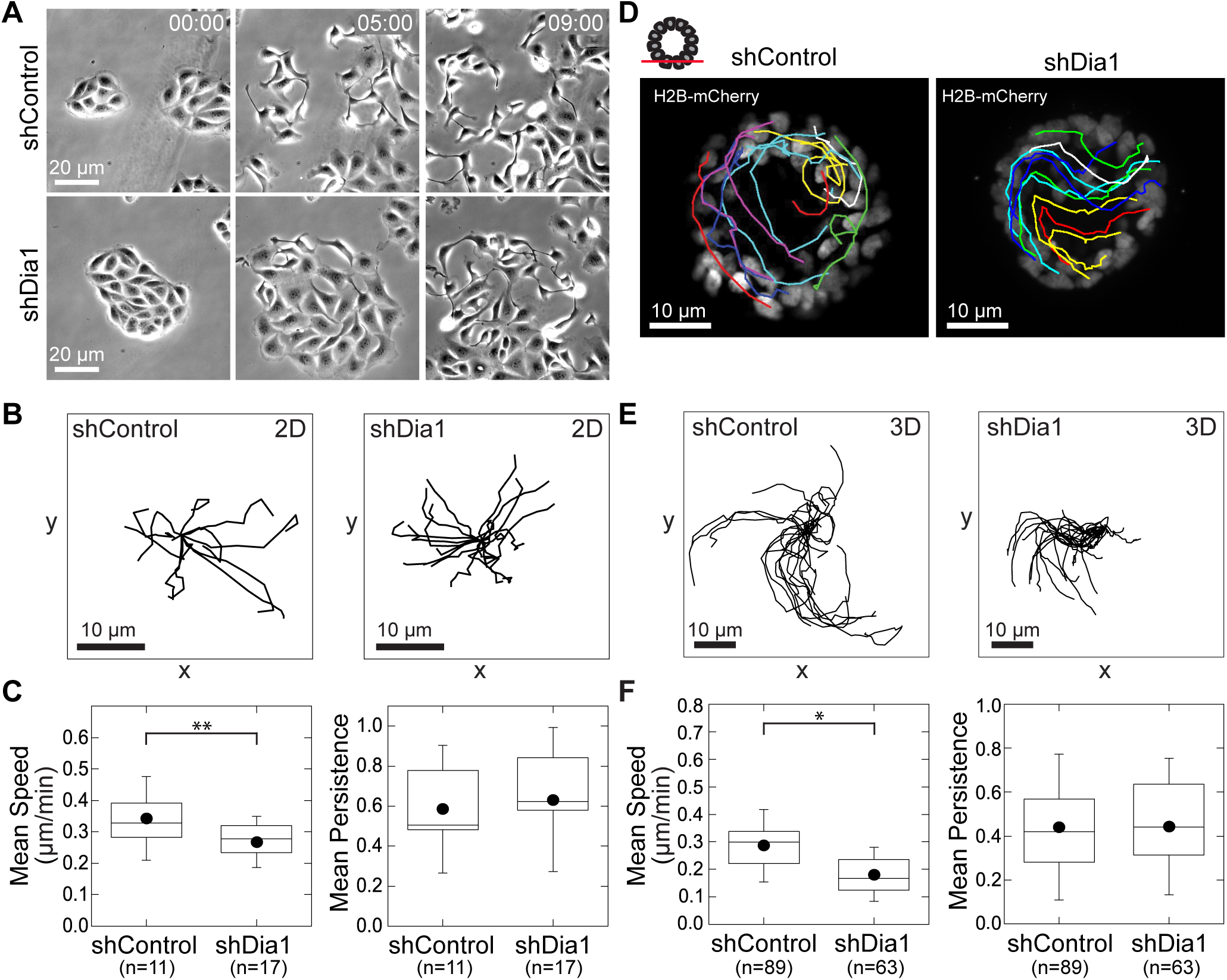
Dia1 is Dispensable for HGF-mediated Planar Motility in 2D and within Acini. **A**, Brightfield images at 0, 5, and 9 hrs of shControl and shDia1 cell islands scattering following 20ng/ml HGF addition at 0 hrs (see also Video 2). Time is indicated in hr:min. **B**, Rose plots of 10 cell trajectories from the cell islands scattering in Fig. 3A and Video 2 over 9 hr. Initial location of each trajectory was positioned at (0,0). **C**, Box plots of instantaneous speed and persistence for shControl and shDia1 cells. **D**, Acini of shControl and shDia1 expressing H2B-mCherry. Acini were stimulated with 20 ng/ml HGF and imaged by timelapse fluorescence microscopy for 12 hrs (see also Video 3). Shown are stills with individual tracks overlaid. **E**, Rose plots from cell trajectories obtained from Fig. 3D and Video 3. **F**, Box plots of instantaneous speed and persistence to characterize cell motility, with number of cell trajectories obtained from at least 5 acini per condition indicated below. Box plots show the 25^th^ and 75^th^ percentiles and the median, circle indicates mean, and whiskers mark 1.5 standard deviations. Single asterisk indicates p<0.01, double asterisk indicates p<0.05 by a Student's two tailed t-test assuming unequal variance.

We next explored cell motility within acini. Cell rearrangements and rotation within 3D tissues have been reported *in vitro* (Pearson and Hunter, 2007), and can contribute to tissue morphogenesis (Ewald et al., 2008; Tanner et al., 2012) and ECM deposition (Isabella and Horne-Badovinac, 2016; Wang et al., 2013). We speculated that MDCK cells may exhibit similar in-plane motility prior to and, perhaps, during branching morphogenesis. To track cell motility within acini, we generated Dia1 knockdown cell lines and matched controls expressing the nuclear marker H2B-mCherry. Immediately after stimulating with HGF, we imaged acini from these cell lines via time-lapse confocal microscopy. Shortly after the start of imaging, shControl cells began moving within the acinus (Fig. 3D and Video 3). This motility resulted in rotation of the entire acinus, with occasional cell rearrangements and rare single cells moving independently of their neighbors. The cell motility within shDia1 acini was virtually indistinguishable from that of control acini (Figs. 3D). Analysis of individual cell tracks (Fig. 3E) showed a decline in average instantaneous speed of shDia1 cells from 0.3 μm/min to 0.2 μm/min, while average persistence did not change relative to shControl cells (Fig. 3F).

These data show that in-plane cell motility driven by HGF, encompassing both scattering of small islands on 2D substrates and motility within acini, does not require Dia1. This suggests that HGF-mediated motility within the plane of a tissue or on 2D substrata is largely independent of Dia1. We next explored whether Dia1 regulates protrusions extending from acini into the surrounding collagen matrix.

### Dia1 is required to stabilize protrusions into the collagen matrix

Defects in branching morphogenesis rendered by Dia1 depletion could arise from reduced formation of protrusions into the collagen matrix. Alternatively, reduced focal adhesion maturation could impair protrusion stability due to reduced adhesion to collagen fibrils.

To adjudicate between these hypotheses, we turned to time-lapse imaging in transmitted light and analyzed protrusive activity at the onset of branching morphogenesis. Following HGF stimulation, shControl acini extended narrow protrusions into the collagen gel over a period of tens of minutes (Fig. 4A and Video 4). All shControl acini imaged formed protrusions over 12 hours of imaging (Fig. 4B). We analyzed protrusion lifetimes by measuring the duration over which protrusions remained before they underwent retraction. This analysis revealed that 60% of all protrusions retracted partially or completely within one hour (Fig. 4C). A smaller proportion were stable for up to two hours, and 20% of protrusions were stable over several hours. These long-lived protrusions eventually mediated cell egress from control acini into the collagen matrix (Video 4). Comparing the lifetime of each protrusion to its maximum length confirmed that the subpopulation of most stable protrusions grew to the greatest lengths of >20 microns (Fig 4D).

**Figure 4:**
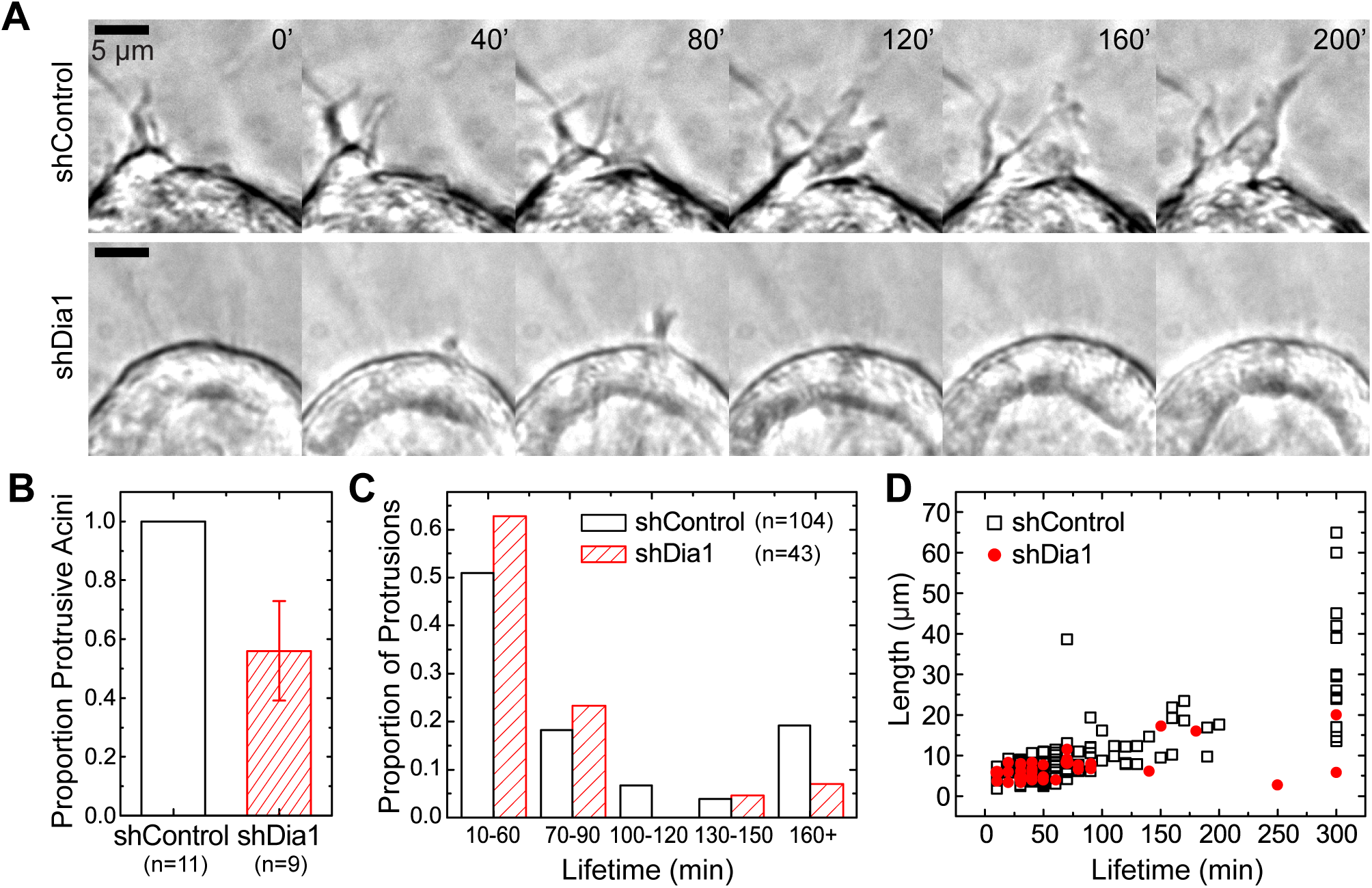
Dia1 is Required to Stabilize Protrusions during Branching Morphogenesis. **A,** Timelapse imaging in brightfield during initiation of branching morphogenesis, showing formation and growth of protrusions in shControl and shDia1 acini over a period of 200 min. Images were obtained starting 3 hrs after addition of 20 ng/ml HGF (see also Video 4). **B,** The proportion of acini from each cell type that formed any protrusions over 12 hrs of imaging, with number of acini scored indicated below. **C,** Histogram summarizing the distribution of protrusion lifetimes from acini for each cell type, from movies obtained 1 hr after HGF addition. Number of protrusions scored indicated in legend. **D,** Plot of protrusion lifetime as a function of its maximal length for protrusions obtained in Fig. 4C.

In contrast to control acini, shDia1 acini exhibited impaired protrusion stability (Fig 4A and Video 4). While 50% of shDia1 acini formed protrusions, the distribution of protrusion lifetimes was weighted towards short protrusions lasting less than 90 minutes (Fig. 4B and 4C). The loss of stable protrusions lasting >2 hours in shDia1 acini was matched by a reduction in the maximum length they reached (Fig. 4D). Meanwhile, protrusions lasting less than 90 minutes were indistinguishable between shDia1 and shControl acini (Fig. S4A). To test whether this failure of protrusion stability caused motility defects that were cell-autonomous, we performed time-lapse imaging of single MDCK cells in collagen gels. This revealed a significant decrease in the motility of single shDia1 cells relative to their shControl counterparts following stimulation with HGF (Fig. S4B and S4C, and Video 5).

These data show that, under normal conditions, the early stages of branching morphogenesis are characterized by numerous small protrusions extending from the acinus into the surrounding collagen matrix. While a majority of these protrusions are unstable and retract within 90 minutes, a minority of protrusions adhere stably and extend into the collagen matrix. The correlation observed between protrusion lifetime and maximum length indicates that protrusion growth depends acutely on their ability to adhere stably to collagen. Dia1 knockdown did not abolish protrusions, nor did it alter lifetimes and sizes of the short-lived protrusions relative to controls. Rather, Dia1 was required for protrusions to stably adhere to collagen matrix and subsequently elongate. These data therefore strongly suggest that Dia1 conditions branching morphogenesis through its role in focal adhesion maturation and stability.

### Dia1 is required to adhere to and displace individual collagen fibrils

To test whether unstable protrusions observed in shDia1 acini resulted from an underlying adhesion defect, we used fluorescence live cell imaging to examine actin, myosin and collagen fibrils during cell protrusion. We imaged shControl and shDia1 acini coexpressing GFP-Lifeact and mCherry-tagged myosin light chain (MLC) plated in collagen matrices labeled with Alexa-647. After incubating acini for 4 hours in HGF, we captured fluorescence confocal images at the lower surface of acini juxtaposed to the collagen matrix. To better capture dynamic cell adhesions to collagen, we acquired images at 3 minute intervals for 3 hours. We noted passive responses of the collagen matrix as cells moved within acini. However we also observed shControl cells deforming single collagen fibrils at discrete points with displacements of ∼2 μm while leaving surrounding fibrils undisturbed (Fig. 5A and Video 6). We captured deformations within each field of view and found an average rate of one per hour per acinus (Fig. 5C). Interestingly, we observed dense MLC puncta and increased actin intensity at sites of fibril deformation, which tracked the collagen fibril as it deformed over tens of minutes (Fig. 5A and Movie 6).

**Figure 5:**
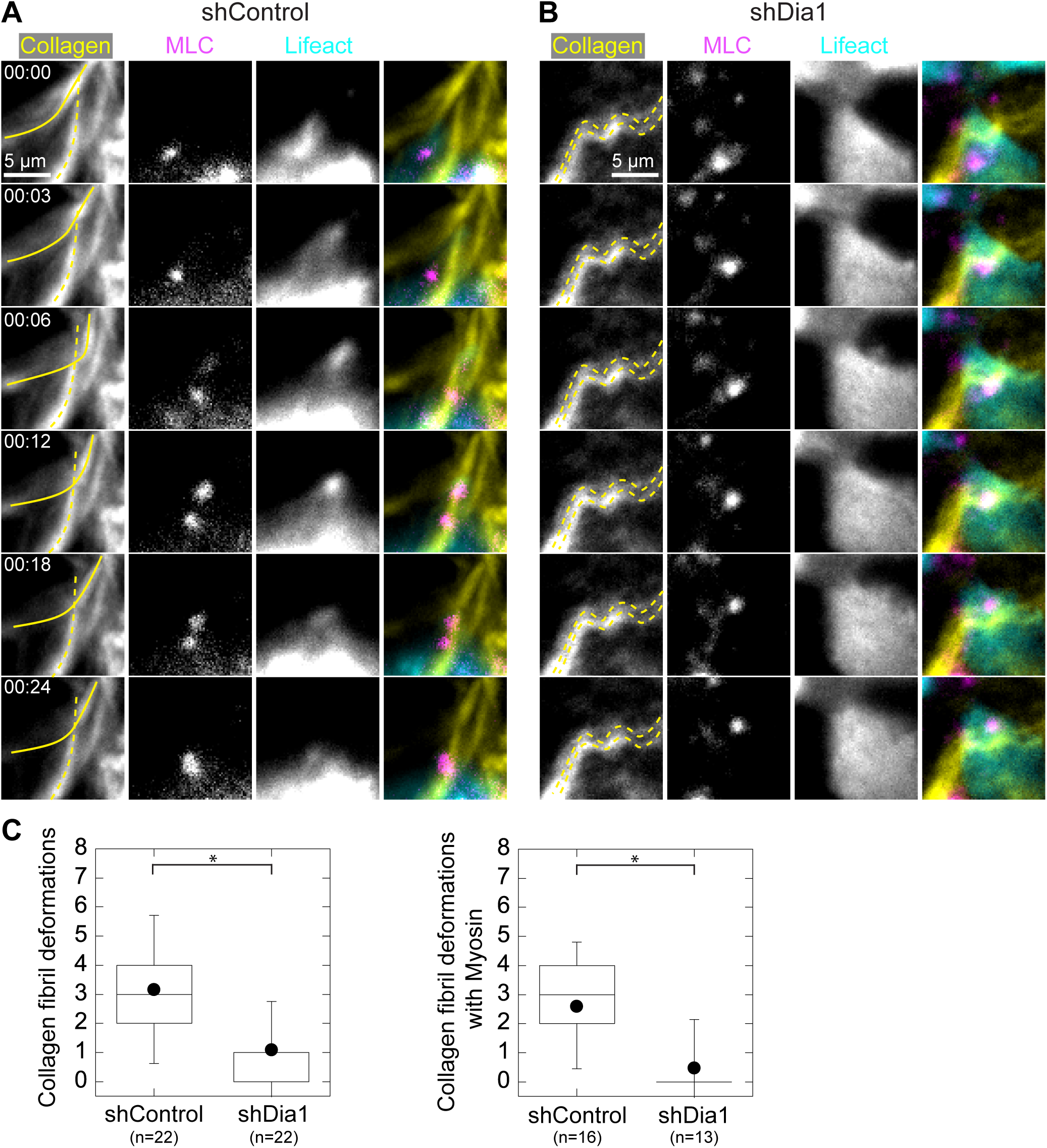
Dia1 is Required to Acutely Displace Individual Collagen Fibrils. **A** and **B**, Images at the basal acinar surface of GFP-Lifeact (cyan), mCherry-Myosin Light Chain (MLC, magenta) and Alexa-647 labelled collagen (yellow) obtained 4 hrs after addition of 20ng/ml Hepatocyte Growth Factor (HGF) in (A) shControl acini and (B) shDia1 acini (see also Videos 6 and 7). Montage in (A) shows an example of shControl cell deforming a single collagen fibril, outlined in solid yellow line, over a period of 24 min. An unaffected fibril is outlined with a dashed line. **B,** Example of shDia1 cell showing collagen fibrils that remain immobile as a cell moves over them. **C,** Box plot indicating frequency of collagen fibril deformations, with number of acini scored indicated below. **D**, Collagen deformations at which MLC appeared or was recruited. Box plots show the 25^th^ and 75^th^ percentiles and the median, circle indicates mean, and whiskers mark 1.5 standard deviations. Asterisk indicates p<0.01 by a Student's two tailed t-test assuming unequal variance.

We observed substantially fewer instances of acute collagen deformations in shDia1 acini, although nonspecific movements of the collagen matrix occurred as with controls (Fig. 5B and Movie 7). On average, we observed a three-fold decrease in collagen fibril deformations by shDia1 acini relative to controls (Fig. 5C, left). When we scored only those deformations with coincident MLC puncta, these were even more suppressed in shDia1 acini relative to controls (Fig. 5C, right).

These data identify Dia1-dependent interactions between cells and the collagen matrix during the earliest stages of HGF-stimulated protrusion. Adhesions to collagen are marked by dense myosin puncta through which cells deform single collagen fibrils while leaving surrounding fibrils unaffected. The planar motility observed in shDia1 acini suggests that Dia1 is not necessary for weak adhesion to collagen fibrils that can support planar motility and rotation, but is required specifically for adhesion to and contraction against collagen fibrils during the onset of branching morphogenesis.

### Myosin co-localizes with Dia1-dependent focal adhesions

The observation that MLC accumulates into discrete puncta at sites of adhesion to collagen suggested that myosin-rich puncta are sites of focal adhesions. To confirm this, we imaged actin, myosin and collagen as leader cells protruded away from acini into the surrounding collagen 24 hours after stimulation with HGF (Video 8). Leader cells advanced via bursts of actin, followed by the accumulation of MLC in puncta at the cell front (Fig. 6A). MLC puncta at the cell front coincided with points of collagen fibril deformation.

**Figure 6:**
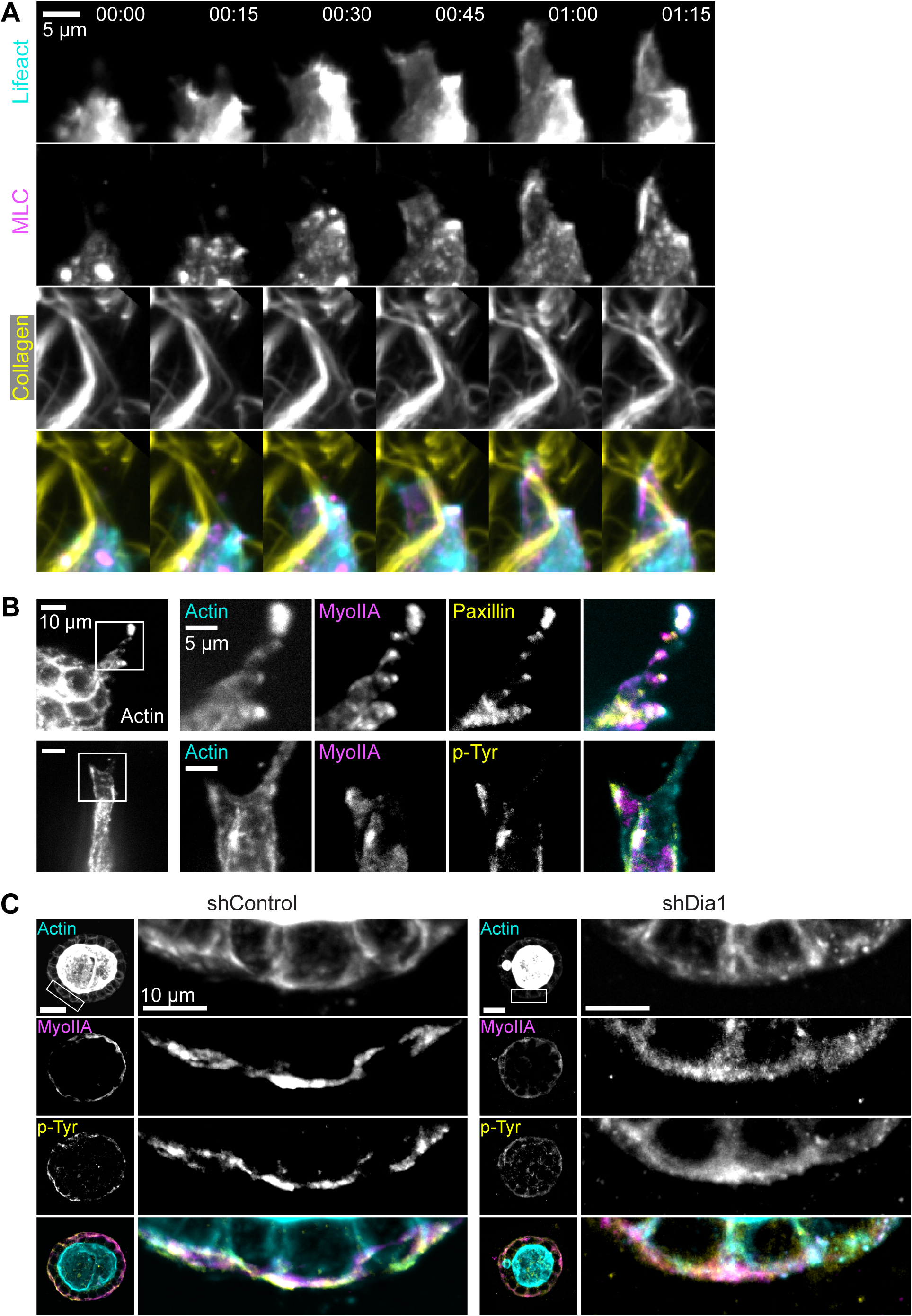
Myosin Co-localizes with Dia1-dependent Focal Adhesions. **A,** Timelapse images of a leader cell protruding into the surrounding collagen matrix from an shControl acinus stimulated with HGF for 24 hrs. Images of GFP-Lifeact (cyan), mCherry-MLC (magenta) and Alexa-647 labelled collagen (yellow) are shown (see also Video 8). Time is indicated in hrs:min. **B,** Immunofluorescence images of f-actin (cyan), Myosin IIA (magenta) and phospho-tyrosine (top row) or paxillin (bottom row), shown in yellow, in leader cells extending from shControl acini following stimulation with HGF for 48 hrs (see also Fig. S5A). **C,** Immunofluorescence images of f-actin (cyan), Myosin IIA (magenta) and phospho-tyrosine acini 4 hrs following stimulation with HGF, combined into maximum intensity projections spanning 3μm. Insets of the indicated boxed regions are shown at right. Scale bars, 10 μm (see also Fig. S5B-C).

To confirm these myosin-rich sites contained canonical focal adhesion proteins, we performed immunofluorescence staining for actin, myosin IIA, and either paxillin or phospho-tyrosine. These images revealed that actin and myosin-rich puncta within leader cells also contained paxillin and phospho-tyrosine (Fig. 6B). We found that around half (0.48 ±0.16, n=283) of all phospho-tyrosine positive puncta were also enriched in myosin IIA (Fig. S5A). These observations confirm that focal adhesions along collagen fibrils retain a punctuate appearance and a subpopulation of them are rich in actin and myosin. While the myosin localization we observed contrasts with canonical descriptions of myosin exclusion from focal adhesions on planar 2D surfaces (Vicente-Manzanares et al., 2009), our observations agree with reports identifying myosin recruitment at leading edge adhesions in 2D and 3D contexts (Pasapera et al., 2015; Wyckoff et al., 2006).

We next assessed the impact of Dia1 depletion on adhesion formation. Because shDia1 acini failed to form stable protrusions into the collagen matrix, we analyzed acini after 4 hours of HGF stimulation, when acinar rotation and collagen interactions can be observed. Acini were fixed and immunostained for actin, phospho-tyrosine and myosin IIA, and maximum intensity projections at the equatorial plane were analyzed. Phospho-tyrosine was enriched in dense puncta at the basal surface in shControl acini (Fig. 6C, left). Meanwhile, phospho-tyrosine was diffusely distributed at the basal surface of shDia1 acini (Fig. 6C, right), and organized into fewer discrete puncta per acinus (Fig. S5B). This defect was accompanied by a redistribution of phospho-tyrosine away from the basal surface in shDia1 acini (Fig S5C), which was paralleled by myosin IIA and paxillin localization (Fig. S5D). We confirmed these results by immunofluorescence staining for mature focal adhesions in cells scattering on glass coverslips. Consistent with prior reports (Oakes et al., 2012), shDia1 cells formed significantly fewer phospho-FAK-positive focal adhesions relative to shControl cells (Fig. S5E and S5F).

These data demonstrate that a subset of adhesions to collagen fibrils are marked by actin and myosin-rich puncta. Myosin accumulation correlates with force generation against collagen fibrils. We find these adhesions do not form in the absence of Dia1, preventing stabilization of protrusive fronts into the collagen matrix. Dia1 has been previously shown to be essential for focal adhesion maturation on 2D surfaces (Oakes et al., 2012), and we suspect that Dia1 plays a similar role here. Clearly, because Dia1 is not required for planar acini motility, adhesions to collagen are capable of forming, consistent with prior observations (Oakes et al., 2012). Rather, we hypothesize that Dia1-mediated focal adhesion maturation is necessary to form stable adhesions that facilitate the egress of cells away from the acini into the surrounding collagen.

### Dia1 regulates actomyosin organization and dynamics of the basal cortex

Prior reports have demonstrated that formin-dependent stress fibers act as templates for the compositional and morphological maturation of focal adhesions (Dolat et al., 2014; Iskratsch et al., 2013; Oakes et al., 2012). We hypothesized that Dia1 performs a similar role at the basal actin cortex of acini, and may control focal adhesion maturation through its effects on actin architecture. Indeed, confocal sections of acini stained with phalloidin revealed striking differences in the organization of cortical actin between shControl and shDia1 acini. In the absence of HGF, cortical actin in shControl acini was organized into bundles often emanating from dense puncta in the middle of cells (Fig. 7A, left). In shDia1 acini, the basal cortex was largely absent of such actin bundles and their associated puncta (Fig. 7A, right). Instead, actin was diffusely spread across the cortex. These differences in basal actin organization remained after HGF stimulation, and were matched by altered myosin localization (Fig. 7B). This confirmed an important role for Dia1 in regulating the organization of the basal actomyosin cortex of acini.

**Figure 7:**
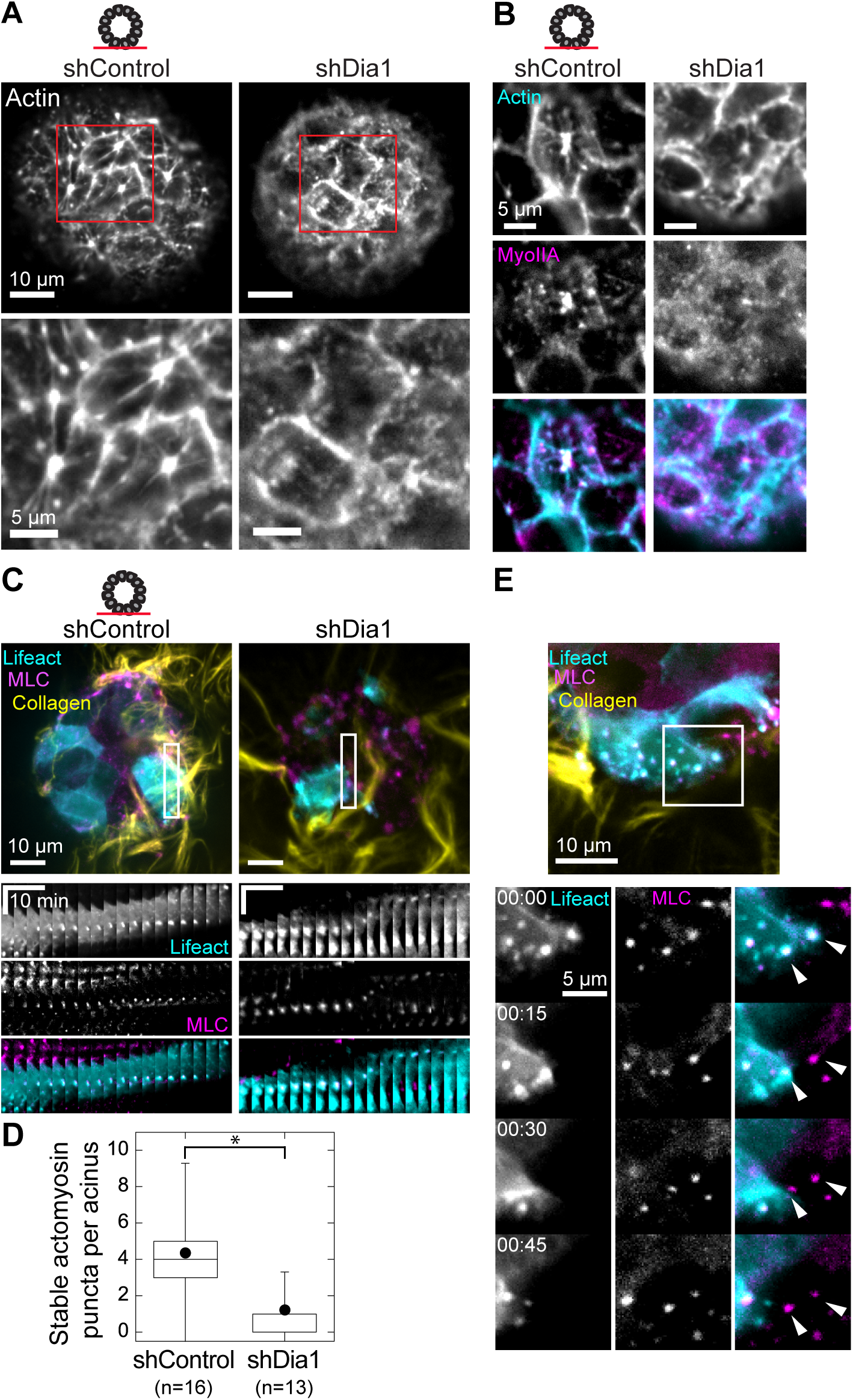
Dia1 Regulates Actomyosin Organization and Dynamics of the Basal Cortex. **A**, F-actin stain of the basal surface of acini prior to HGF addition, with region of interest enlarged below. **B**, Immunofluorescence of the basal surface of f-actin (cyan) and Myosin IIA (magenta) in shControl and shDia1 acini stimulated with 20ng/ml HGF for 4 hrs. **C**, shControl and shDia1 acini coexpressing GFP-Lifeact (cyan) and mCherry-MLC (magenta) were stimulated for 4 hrs with 20ng/ml HGF and fluorescence images were taken every 3 min at the basal surface (see also Video 9). Kymographs show representative MLC and actin puncta, time scale bar represents 10 min and distance scale is 10 μm. **D**, Box plot of the number of stable actomyosin puncta per field of view observed over 3 hr window, with total organoids imaged indicated below. **E**, Images of GFP-Lifeact (cyan) and mCherry-MLC (magenta) and Alexa-647 labelled collagen (yellow) in shControl acinus (see also Video 10). Montage shows movement of a cell with high expression of GFP-Lifeact, followed by a low-expressing cell. As the first cell moves to the left, MLC and Lifeact puncta remain stationary (arrowheads). Once the cell moves past this region, the next cell forms puncta at the same location. Box plot shows the 25^th^ and 75^th^ percentiles and the median, circle indicates mean, and whiskers mark 1.5 standard deviations. Asterisk indicates p<0.01 by a Student's two tailed t-test assuming unequal variance.

To explore changes in dynamics of the actomoysin cortex, we used timelapse fluorescence microcroscopy to image the basal surface of GFP-Lifact/MLC-mCherry acini (Fig. 7C). We found MLC puncta in shControl remained immobile in cells moving within the plane of the acinus over tens of minutes (Fig. 7C and Movie 9). By contrast, MLC puncta in shDia1 were more dynamic, undergoing assembly and motion over similar time intervals (Fig. 7C and Movie 9). Using a threshold of 12 minutes without displacement to distinguish immobile from mobile MLC puncta, we found that shControl acini formed three-fold more immobile MLC puncta relative to shDia1 acini (Fig. 7D).

How do individual cells adhere to and migrate along collagen fibrils in the midst of ongoing planar motility within the acinus? During acinar rotation, we observed isolated instances of multiple cells forming MLC puncta at the same location on collagen fibrils as they encountered it in sequence (Fig. 7E and movie 10). Interestingly, the next cell encountering the same region of collagen assembled a puncta in the same location. This example illustrates that certain locations within the collagen matrix are primed for repeated adhesion assembly in subsequent cells during planar acinar motility.

## Discussion

The suite of molecular mechanisms governing cell motility through 3D environments has enjoyed much attention, as these form the basis for profound tissue shape changes during development and cancer invasion (Gray et al., 2010). Detailed models exist to describe how cytoskeletal organelles control the motility of single cells (Petrie and Yamada, 2012), but how these organelles are regulated in space and time to effect tissue shape changes is less clear. Data presented here suggest a previously unappreciated mechanism by which focal adhesion maturation regulates branching morphogenesis in a model epithelial tissue.

Our results suggest that focal adhesions in MDCK acini embedded in collagen matrices share key features with focal adhesions in cells on 2D substrata. We and others have previously reported that maturation of focal adhesions during 2D cell motility requires a stress fiber template, through the activities of alpha-actinin, septins and formins such as Dia1 (Choi et al., 2008; Dolat et al., 2014; Gupton et al., 2007; Oakes et al., 2012). HGF signaling in MDCK cells activates Rho, which regulates Dia1 and myosin activity (Ridley et al., 1995). These effectors play critical roles during branching morphogenesis. When an MDCK cell initially contacts a collagen fibril, formation of a nascent adhesion causes a burst of actin polymerization and myosin recruitment (Fig. 8A). Our data indicate that actin and myosin concentrate at the adhesion in a Dia1-dependent manner and promote its maturation, marked by phospho-tyrosine rich proteins such as Focal Adhesion Kinase (FAK) and Paxillin (Mitra et al., 2005; Zaidel-Bar et al., 2006). Mature adhesions are resistant to disassembly as cells change shape and move within the acinus. In concert with myosin contractility, mature focal adhesions enable cells to pull themselves and their neighbors away from the acinus into the collagen matrix (Fig. 8A).

**Figure 8:**
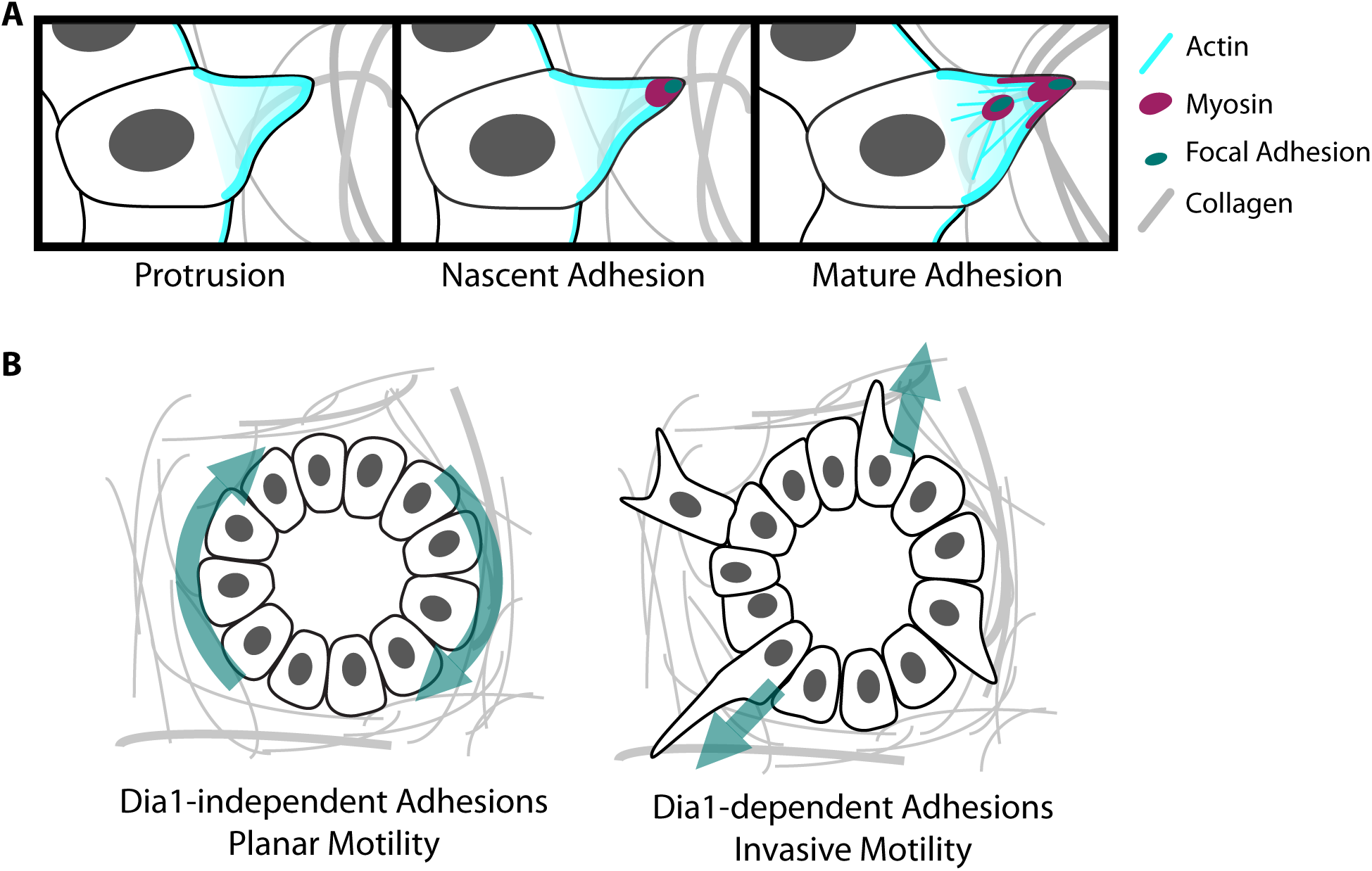
Maturation of focal adhesions through Dia1 during tissue shape changes. **A,** Model for focal adhesion maturation during the onset of MDCK branching morphogenesis. Cells generate actin-rich protrusions from their basal surface into the collagen matrix. Protrusions are initially weakly adhered to collagen fibrils through nascent focal adhesions. Localized actin polymerization through Dia1 and myosin recruitment stabilize focal adhesions and promote their maturation. Mature adhesions resist turnover and allow cells to exert contractile forces against collagen fibrils to enable branching morphogenesis. **B,** Dia1 may function as a mechanism by which tissues can regulate noninvasive and invasive motility. In the absence of Dia1 activity cells can adhere sufficiently to the collagen matrix to mediate planar motility within the acinus and acinar rotation. Invasive motility into the collagen matrix requires that cells form mature adhesions, dependent on Dia1.

This work provides evidence to expand the biological functions of focal adhesion maturation. Despite the exquisite detail with which focal adhesion assembly and maturation has been defined (Geiger and Yamada, 2011), the acute consequences of maturation defects on cell motility and tissue morphogenesis remain poorly defined (Fraley et al., 2010; Hoffman et al., 2006; Sieg et al., 2000; Thievessen et al., 2013). Data presented here suggest that tissues may require focal adhesion maturation to carry out complex shape changes in 3D fibrillar environments. This conclusion is supported by a recent report demonstrating that focal adhesion maturation through septins is similarly required for MDCK branching morphogenesis (Dolat et al., 2014). Our data showing a similar role of formins during migration by mouse tumor explants into collagen matrices suggests this idea could be generalized to other morphogenic processes. For an individual cell within a developing or malignant tissue, it is not clear *a priori* how its motility within the tissue or into the ECM is determined. Here we present a mechanism by which adhesions to the ECM act as a switch from planar motility to invasive motility (Fig. 8B). Together, these results suggest an expanded repertoire for the biological functions of focal adhesions and their regulators, and prompt testable predictions about the interplay between adhesion stability, cell motility, and tissue shape.

In contrast to adhesions in 2D contexts, we observed myosin accumulation at sites of adhesion in acini. Our data suggest that adhesions to collagen fibrils are sites of continuous actin polymerization and sustained myosin recruitment. It is possible that Rho is recruited to adhesions, and mediates local Dia1 and myosin activation. Indeed this role is consistent with observations of Rho at the leading edge of migrating cells (Fritz et al., 2013; Machacek et al., 2009). Finally, the recruitment of myosin at adhesions is a feature shared with invadosomes (Collin et al., 2008; Parekh and Weaver, 2016). Formation of invadosomes in cancer cells were shown to require formins (Lizarraga et al., 2009), raising the possibility that adhesions described here also play proteolytic roles during MDCK branching morphogenesis.

The role we identify for Dia1 carries important implications for its primary regulator, Rho. Rho activation is thought to impact cell motility primarily through ROCK and myosin contractility (Petrie and Yamada, 2012). Prior studies have shown that reducing myosin activity during branching morphogenesis results in increased branching, implicating Rho as a negative regulator of tissue shape changes (Fischer et al., 2009; Yu et al., 2003). Conversely, we show that Dia1 is required for branching morphogenesis through stabilizing focal adhesions to collagen fibrils. Together these observations suggest competing effects of Rho signaling during complex tissue shape changes: one as a negative regulator via ROCK-mediated contractility and one as a positive-regulator via Dia1-mediated focal adhesion maturation. In the absence of ROCK-mediated contractility, focal adhesion maturation is not needed to facilitate migration of cells away from the acinus due to the overall low levels of contractile tension within the tissue. In the presence of ROCK-mediated contractility, Dia1-mediated focal adhesion maturation is not required for weak adhesion to collagen fibrils and planar cell motility to support rotation. However, it is required to assemble adhesions that are sufficiently stable to withstand physiological contractile tension as cells migrate away from the acinus and drive tissue shape change. Future studies may delineate these apparent opposing roles of Rho effectors, and whether they indeed converge on focal adhesion stability and spatial regulation of contractility during tissue morphogenesis.

## Materials and Methods

### Cell culture and reagents

Manin-Darby Canine Kidney (MDCK) type II/G cells (American Type Culture Collection) were cultured in a humidified incubator with 5% CO_2_ using Dulbecco’s Minimal Essential Medium (Corning) supplemented with 5% fetal bovine serum (FBS) (Corning), 2 mM l-Glutamine, and penicillin-streptomycin (Corning). Selection media was supplemented with 5 μg/ml puromycin (Gibco). The MDCK cell line expressing GFP-Lifeact was a gift from Thorsten Wittman (University of California, San Francisco). The following antibodies were used: Rat anti-Ecadherin DECMA (Santa Cruz Biotechnologies), mouse anti-Podocalyxin (gift from the Keith Mostov Laboratory, University of California, San Francisco), rabbit anti-Laminin 511 (Sigma), rabbit anti-Myosin IIA heavy chain (Covance), mouse anti-Paxillin 5H11 (Millipore), rabbit anti-FAK-pY397 (Life Technologies), Alexa fluor 488 goat anti-mouse secondary (Life Technologies), Alexa fluor 568 goat anti-mouse secondary (Life Technologies), Alexa fluor 568 goat anti-rat secondary (Life Technologies), rabbit anti-Diaph1 (ProteinTech), rabbit anti-Diaph2 (Cell Signaling), rabbit anti-FHOD1 (Abcam), rabbit anti-GAPDH. For immunofluorescence, all primary antibodies were used at 1:200 and all secondary antibodies were used at 1:500. The formin inhibitor SMIFH2 (gift from David Kovar, University of Chicago) was dissolved in DMSO and used at 30 μM. The Arp2/3 inhibitor (Calbiochem) was dissolved in DMSO and used at 50 μM.

### DNA constructs and gene knockdown

Knockdown cell lines were generated using shRNA constructs against canine Dia1 (gatccccgccacagatgagagagacattcaagagatgtctctctcatctgtggcttttta), Dia2 (gatccccgcaaccttacagcaatggattcaagagatccattgctgtaaggttgcttttta), FHOD1 (gatccccgaagacgaggacatactgattcaagagatcagtatgtcctcgtcttcttttta), or a nontargeting control (OligoEngine). These were annealed and cloned into the HindIIII and BglII sites of the pSuper retroviral vector (OligoEngine). Retrovirus was produced using the Phoenix cell line (Nolan Lab, Stanford University) using Fugene 6 transfection reagent (Roche) to transfect the retroviral vector and a VSV-G psuedotyping plasmid (gift from Marsha Rosner, University of Chicago). Viral supernatant was collected, filtered, and incubated with target MDCK cells for 12 hours in the presence of 8 ug/ml polybrene (Millipore).Following viral transfection and selection, knockdown was confirmed by western blot. For nuclear tracking, shControl and shDia1 cells were transfected using a pQCXIX retrovirus construct encoding H2B-mCherry (gift from Mark Burkard, University of Wisconsin). Cells were purified using flow cytometry (University of Chicago Flow Cytometry Core). The lentiviral vector encoding MLC-mCherry was generated by cloning MLC-mCherry sequence into pWPT lentiviral backbone (Addgene plasmid 12255) with the aid of SnapGene Software (GSL Biotech LLC; www.snapgene.com). Virus was produced in 293T cells (gift from Geof Green, University of Chicago) using a pHR1-8.2-deltaR packaging plasmid and a VSV-G psuedotyping plasmid (gifts from Marsha Rosner, University of Chicago). Following viral transfection cells were isolated by flow cytometry at the University of Chicago Flow Cytometry Core to purify mCherry and GFP positive cells.

### Culture and manipulation of acini

Matrigel (Corning) from a single lot containing 9.1mg/ml protein was used for all 3D culture experiments. To generate acini, the lower surface of wells in a 24-well plate were coated with 30 μl Matrigel and allowed to gel at 37° C. MDCK cells were trypsinized, pipetted vigorously to break up cell clusters, and 10,000 cells were added to 800 μl of a 1:1 solution of growth media and Matrigel, which was kept on ice to prevent gelation. The resulting cell suspension was immediately pipetting up and down to disperse cells and added to the Matrigel-coated well The Matrigel-cell suspension was immediately placed in a 37° C incubator and allowed to gel for 30 minutes before 800 μl growth media was gently added to each well. Media was changed every 3 days for 6-7 days until acini formed lumens. To isolate acini, a modified protocol by Rubashkin et al(2014) was used to melt Matrigel by incubation at low temperatures. Briefly, media was aspirated and Matrigel was disrupted by pipetting up and down with 5 ml of warm PBS with Ca/Mg (Corning) per well. The PBS-Matrigel solution was pelleted at 500 x g for 3 minutes and resuspended in 10 ml fresh PBS was added. Acini were incubated in a bucket of salted ice with rocking at maximum speed for 40 minutes. Acini were pelleted at 500 x g, yielding a ∼50 μl slurry of residual Matrigel and acini which were kept on ice to prevent gelation. Collagen gels were prepared 30 minutes before use and kept on ice. To prepare collagen gels, 1 M HEPES and 7.5% NaHCO3 were combined with media to achieve final ratios of 1:50 and 1:23.5, respectively. Rat tail collagen 1 (Corning) was gently added to a final concentration of 2 mg/ml. Collagen was fluorescently labeled using Alexa Fluor 647 NHS ester (Life Technologies), dissolved at 10 μg/μl in DMSO and incubated with Rat tail collagen at a ratio of 1:1000 and stored at 4 C. The resulting Alexa-647-labelled collagen was included in a ratio of 1:4 with unlabelled collagen. Acini were added to the collagen solution, mixed by pipetting, and plated in either 4 or 8-well Ibidi chambers (Ibidi) or in 4-well Labtek chamber slides (Nunc), in volumes of 40 or 100 μl. All chambers were precoated with 10 or 30 μl collagen solution for 10 minutes prior to plating acini. Collagen was allowed to gel for at least 30 minutes before growth media was gently added to the sides of wells.

### Mouse tumor explants

One female day 70 mouse of the Mouse Mammary Tumor Virus-Polyoma Middle T Antigen strain (MMTV-PyMT, Jackson Laboratories) was sacrificed and the largest mammary tumor from each inguinal mammary gland was surgically excised. Each tumor was manually minced with a razorblade and tumor tissue was digested by shaking for 30 minutes at 37 C in a conical tube containing 40 ml of DMEM/F12/50:50 (Corning) with 3 mg/ml Collagenase A, 1 mg/ml hyaluronidase (Worthington) and 2 U/ml DNase 1 (Fisher). 5 ml of FBS was added to halt digestion, and tumor tissue suspension was pelleted at 500x g for 3 minutes and resuspended. Tumor organoids were placed in a 1:1 mixture of Matrigel and DMEM/F12/50:50 supplemented with 10% Fetal Bovine Serum (Corning), 10 μg/ml human Insulin (PromoCell), 5 μg/ml human Transferrin, and penecilin/streptomycin (Corning). The bottom wells of a 24-well plate (Corning) were coated with Matrigel and allowed to gel before adding 800 μl of organoid-Matrigel suspension per well. Matrigel was allowed to gel for 30 minutes and 800 μl growth media was gently added. Organoids were cultured in Matrigel for 24-48 hours prior to replating into 2mg/ml Collagen 1 gels in 4-well Ibidi chambers as described above, with the ice incubation step reduced to 20 minutes. Following replating in collagen, fresh media containing DMSO, 30 μM SMIFH2, or 50 μM CK666 was added and chambers were placed on a heated microscope stage. Images were collected every 30 minutes for 72 hours.

### Western blot analysis

For Western blotting, cells were lysed in Laemmli buffer (4% sodium dodecyl sulfate, 20% glycerol, 120 mM Tris-Cl pH 6.8, 0.02% bromophenol blue). Lysates were separated by SDS-PAGE gel and electrotransferred to a nitrocellulose membrane. Blots were blocked in PBS with 5% nonfat dry milk and incubated with primary antibodies at 1:1000 overnight at 4 C. Blots were incubated in secondary antibodies at 1:10000 for 1 hour at room temperature and developed with ECL Western blotting substrate (Thermo Fisher Scientific). Blots were scanned as film negatives on a photo scanner (Perfection v700; Epson) and analyzed using the gel analysis tool in ImageJ (National Institutes of Health). The intensity of the protein bands was normalized through comparison with the loading control bands.

### Microscopy

All fluorescence images were acquired on an inverted microscope (Ti-E; Nikon) with a confocal scanhead (CSUX; Yokogawa Electric Corporation), laser merge module containing 491, 561, and 642 laser lines (Spectral Applied Research), a stage controller (Prior), and a cooled charge-coupled device camera (HQ2; Roper Scientific). Images were acquired using either a 20x 0.75 NA Plan Fluor multi-immersion objective (Nikon) or a 40x 1.15 NA Plan Apo water immersion extra long working distance objective (Nikon). Immunostained acini or organoids were imaged by acquiring z-stacks with either 500 nm or 1 μm z spacing. All transmitted light images were acquired on an inverted microscope (Ti-E; Nikon), a stage controller (Prior), and a cooled charge-coupled device camera (HQ2; Roper Scientific). Images were acquired using a 20x 0.45 NA air extra long working distance objective (Nikon). All hardware was controlled via MetaMorph acquisition software (Molecular Devices). Brightfield images of acini were acquired on a Nikon Eclipse TS100 (Nikon) with an iPhone 6S (Apple) and a SnapZoom adaptor (SnapZoom).

### Live cell imaging

All live cell imaging was performed with a stage incubator for temperature, humidity, and CO_2_ control (Chamlide TC and FC-5N; Quorum Technologies). The stage adaptor, stage cover, and objective were maintained at 37° C, while humidified 5% CO_2_ air was maintained at 50° C at its source to prevent condensation within its tubing. Acini were transferred to collagen gels in 4- or 8-well plastic chambers (Ibidi) at least 1 day prior to imaging. Imaging media was identical to growth media except phenol red-free DMEM was used. Imaging media containing 20 ng/ml HGF was added to chambers immediately before transfer to the pre-warmed microscope stage, where they equilibrated for ∼30 minutes prior to imaging. Image sequences were corrected for drift using the Stackreg plugin in ImageJ (National Institutes of Health), and a custom Matlab script. Fluorescence imaging of cells expressing H2B-mCherry was performed by collecting z-stacks of 21 planes at 3 μm intervals every 10 minutes. For fluorescence imaging of GFP-Lifeact, mCherry-MLC, and Alexa Fluor 647-collagen, 20 ng/ml HGF was added to phenol red-free media 4 or 24 hours prior to imaging. For each acinus, images were acquired at 6 planes separated by 3 μm every 3 minutes for 3 hours. Only the lower surface of each acinus was imaged, to capture collagen fibrils and cells simultaneously. Alexa Fluor 647-collagen images were bleach corrected in ImageJ.

### Immunofluorescence

For immunofluorescence staining of acini, a 1.5 U/ml solution of Clostridium Collagenase (Sigma) in PBS was added to culture media prior to fixation and acini were replaced in the incubator for 10 minutes. Acini were fixed in a solution of 4% paraformaldehyde with 0.1% Triton X-100 in Phosphate Buffered Saline solution (PBS, Corning). Fixation solution was gently added to each well while simultaneously aspirating culture media, and incubated for 20 minutes. Collagen autofluorescence was quenched with 3 rinses of 0.1 M glycine in PBS for 20 minutes each. Acini were permeabolized in 0.5% Triton X-100 for 10 minutes, and blocked with 2.5% Bovine Serum Albumin and 0.1% Triton X-100 in PBS for 1 hour. Acini were incubated with primary antibody at 1:200 in blocking solution overnight at 4° C, washed 3 times in 0.1% Triton X-100 for 20 minutes each, and secondary antibody at 1:500 or Alexa fluor 647 phalloidin at 1:1000 (Life Technologies) was incubated in blocking solution for 1 hour. After another 3 20 minute washes with 0.1% Triton X-100, chamber walls were removed and 20 μl Prolong Gold (Life Technologies) was added per well. A clean coverslip had clear blobs of nail polish applied to its corners and allowed to dry, and placed over the chamber slide. Slides were allowed to dry, sealed with nail polish, and stored at 4° C. All incubations were performed on a rotary shaker. Immunofluorescence staining of cells in 2D culture was performed as above but without glycine rinses.

### Branching morphogenesis assays

Acini plated in collagen gels were cultured for at least 1 day prior to inducing branching morphogenesis. Briefly, growth media was removed and replaced with media containing 20 ng/ml human recombinant Hepatocyte Growth Factor (HGF) (Sigma-Aldrich), with DMSO or inhibitors as indicated. Fresh HGF with inhibitors was added after 24 hours. After 48 hours, acini were fixed and stained with phalloidin.

### Scattering assays

MDCK cells were plated sparsely in 4-well plastic chambers (Ibidi) or on glass coverslips and allowed to form islands. Cells were serum starved by replacing growth media with phenol red-free media supplemented with 0.5% FBS, and culturing for 12 hours. For timelapse microscopy, chambers were placed on a heated microscope stage and 20 ng/ml HGF was added to low-serum media to stimulate scattering. Cells were imaged in brightfield at 10 minute intervals for 12 hours. For immunofluorescence, cells were plated on glass coverslips and allowed to form islands, then serum starved for 12 hours, incubated in HGF for 6 hours, and fixed for immunofluorescence staining.

### Image Analysis

For branching morphogenesis assays, z-stacks of each acinus were collected at 0.5 or 1 μm intervals and the entire volume was scored for protrusions. These were scored as single- or multi-cellular extensions at least 5 μm in length emanating from acini into the collagen gel. Protrusion length was measured as a straight line from the basal surface of the cell or cells to the tip of the cellular extension. Protrusions extending in the z direction were measured by counting the confocal sections they spanned. For nuclear tracking, image sequences for analysis were generated by combining lower z-planes from each acinus into maximum projections. Cells were tracked manually in ImageJ (National Institutes of Health) and tracks were analyzed using custom Matlab scripts. For cell tracking during scattering, cells were tracked manually using ImageJ (National Institutes of Health) and tracks were analyzed using custom Matlab scripts. Time of initial of cell-cell contact rupture was scored by the first cell to contract and completely dissociate from its neighbors. Quantification of phospho-Tyrosine puncta was performed on maximum intensity projections combining 3 μm at the acinar equator. The resulting images were background subtracted and binarized, and puncta were counted using the Analyze Particles feature of ImageJ (National Institutes of Health), set to a pixel range of 30-1000 pixels^2^. Maximum intensity projections were also analyzed with linescans, each measuring 2 by 15 μm. Quantification of phospho-FAK was performed on background subtracted, binarized images using the Analyze Particles feature of ImageJ (National Institutes of Health), set to a pixel range of 30-1000 pixels^2^. Analysis of collagen fibril deformations and MLC puncta mobility was performed on image sequences in which the lower basal surface of cells and the collagen matrix could be resolved in the same imaging plane. This surface ranged from 1500-2100 μm^2^, representing approximately 0.19 of the total surface area for a typical acinus 50-60 μm in diameter. To quantify collagen fibril deformations, image sequences were analyzed for fibrils that were deformed by 1-2 μm, independently of their neighbors.

### Statistical Tests

To assess statistical significance, we used independent two-sample Student’s *t* tests of the mean to determine the significance with respect to WT or Control. P values were indicated by *, p<0.01; **, p<0.05.

## Acknowledgements

The authors wish to acknowledge with deep gratitude the contributions of Guillermina Ramirez-San Juan and Andrew Ewald for indispensable discussion and guidance, as well as Mark Burkard, Keith Mostov, and Valerie M. Weaver for reagents and protocols. This work was funded by a National Institutes of Health grant (R01 GM104032) to M.L. Gardel, a a National Institutes of Health grant to K.F. MacLeod (RO1 CA162405), and a National Institutes of Health Training Grant (T32 CA00959428).

